# Comparative analysis of varicella-zoster virus and herpes simplex virus 1 interaction with epidermal terminal differentiation in primary human keratinocytes models of differentiation

**DOI:** 10.64898/2026.04.08.717198

**Authors:** Cristina Tommasi, Geonhee Kim, Aoxin Liu, Andriani Drousioti, Olumide Ogunbiyi, Oscar Enrique Torres Montaguth, Afshin Mosahebi, Judith Breuer

**Affiliations:** Infection, Immunity and Inflammation Department, University College London GOS Institute of Child Health, London, UK; Histopathology Department, Camelia Botnar Laboratories, University College London Great Ormond Street Hospital for Children, London, UK; Department of Surgical Biotechnology, Royal Free Hospital, University College London, London, UK

## Abstract

Varicella-zoster virus (VZV) is the etiological agent of chickenpox and herpes zoster, while herpes simplex virus 1 (HSV-1) causes oral and genital herpes. Both infections manifest with skin blisters from which the viruses are transmitted to new hosts either via aerosol (VZV) or skin microabrasions (HSV-1). VZV reaches the skin through the blood route, and in the skin epidermis it first infects undifferentiated keratinocytes of the basal layer. Conflicting evidence exists for HSV-1, making it unclear whether HSV-1 infects undifferentiated or differentiated keratinocytes. Here, we developed *in vitro* models of primary human epidermal keratinocytes’ differentiation to recapitulate infection of distinct layers of the epidermis by VZV and HSV-1. Our data show that replication of both viruses is restricted, VZV more than HSV-1, if initial infection occurs in differentiated keratinocytes, but not if initial infection occurs in basal undifferentiated keratinocytes. Like VZV, HSV-1 downregulates expression of proteins associated with keratinocyte differentiation, such as the suprabasal keratin K10. However, whereas in VZV downregulation of K10 occurs soon after VZV infection and appears to be independent from viral replication, HSV-1-mediated K10 downregulation requires full viral replication. These observations provide insights into the potential for VZV and HSV-1 interactions with epidermal differentiation to yield strategies for developing host- and pathogen-directed antiviral agents.

## Introduction

Varicella-zoster virus (VZV) and herpes simplex virus 1 (HSV-1) are human skin-tropic alphaherpesviruses. Primary infection by VZV causes chickenpox, which presents with a diffuse skin blistering rash [1], while reactivation from neuronal latent state causes herpes zoster (or shingles). HSV-1 is responsible for oral and genital herpes, which present as blistering lesions in the oral/facial and genital skin and mucosa. While VZV reactivation usually occurs once in a lifetime, this is a much more frequent event in infections by HSV-1 [2].

Both VZV and HSV-1 replicate and form lesions in the outermost layer of the skin, the epidermis. The formation of epidermal lesions which contain the infectious virions is essential for transmission to new hosts, as well as for latency as it is likely that the virions encounter nerve endings at the level of the cutaneous blisters [3,4].

The epidermis is a stratified tissue where each stratum is composed of keratinocytes at a specific differentiation state. The basal layer is made up of progenitor (undifferentiated) keratinocytes. A terminal differentiation programme is responsible for gene expression changes in the basal keratinocytes which cause them to leave this layer to populate the suprabasal layers, which are, in ascending order, the spinous, granular, upper-granular and cornified layers. As the keratinocytes move upwards in the tissue, markers of the undifferentiated status, such as keratin 5 (*KRTS*) and *KRT14* are replaced by markers of terminal differentiation. The first markers to be expressed already at the spinous layer include involucrin (*IVL*), keratin 1 (*KRT1*) and keratin 10 (*KRT10*), while markers such as loricrin (*LOR*) and filaggrin (*FLG*) start to be expressed in the granular layer [5,6].

Following haematogenous spread from the nasopharyngeal lymphoid tissue (chickenpox) or reactivation from sensory nerve endings (herpes zoster), VZV infection of skin is first detected in cells within hair follicles from where the infection spreads to interfollicular epidermis basal keratinocytes [7–12]. We previously demonstrated that productive VZV replication and the formation of blisters containing cell-free virions occur only following terminal differentiation of infected basal keratinocytes [13]. In contrast, HSV-1 initially enters the skin through microabrasions, however there is disagreement to whether HSV-1, like VZV, requires to infect basal (undifferentiated) keratinocytes or if infection of suprabasal (differentiated) keratinocytes can also result in productive infection[14–18].

Here, we made use of the calcium switch model [19–21] involving primary human epidermal keratinocytes to investigate more precisely than standard skin models the impact of three specific keratinocyte differentiation states: undifferentiated (basal layer), early differentiated (spinous layer) and late differentiated (granular/upper-granular layer) on VZV and HSV1 infection, replication and life cycle. We show that while both VZV and HSV-1 can infect keratinocytes at all differentiation states, VZV replication is severely restricted in suprabasal differentiated keratinocytes. HSV-1 appears less stringent than VZV with good levels of replication even when infecting differentiated keratinocytes, although replication is highest when undifferentiated basal keratinocytes are infected.

We also demonstrate that HSV-1, like VZV, downregulates keratin 10, a keratin produced only in suprabasally differentiated keratinocytes and which we have previously hypothesised is key to the formation of blisters [13,22]. However, our data suggest differences between the two viruses in the mechanisms by which this occurs.

## Results

### VZV replication and life cycle are inhibited when the virus infects already differentiated keratinocytes

We have previously shown that VZV infects basal progenitor keratinocytes with viral gene expression being tightly linked to keratinocyte differentiation [13]. To determine whether it can also infect the spinous and granular/upper-granular layers, we made use of the calcium switch model [19–21] applied to primary human epidermal keratinocytes, NHEKs. This model enabled us to reconstruct keratinocytes at three distinct stages of epidermal differentiation (**Figure 1A**): undifferentiated (Undiff._**Figure 1B** 1.), early differentiated (Early diff._ **Figure 1B** 2.) and late differentiated (Late diff._ **Figure 1B** 3.). Undifferentiated keratinocytes are characterised by expression of basal layer markers including *KRTS*, early differentiated keratinocytes start expressing the suprabasal markers *KRT1, KRT10* that typify the spinous epidermal layer, but not *LOR, FLG* or *LCE3D* (late cornified envelope 3D), which are only expressed in the late differentiated model representing the granular/upper-granular layers [5,6,23] (**Figure S1**). “Undiff.”, “Early diff.” and “Late diff.” refer to the states of the keratinocytes at the time of infection with VZV (**Figure 1B**_day 0), however, because we previously demonstrated that VZV fully replicates when infected undifferentiated keratinocytes undergo differentiation [13], the “Undiff.” model encompasses the induction of synchronised (through calcium switch) keratinocytes’ differentiation (**Figure 1B** 1._Differentiating_sync.) after infection with VZV. The addition of calcium to cells does not directly affect VZV replication ([13] and **Figure S2A**).

**Figure 1:**
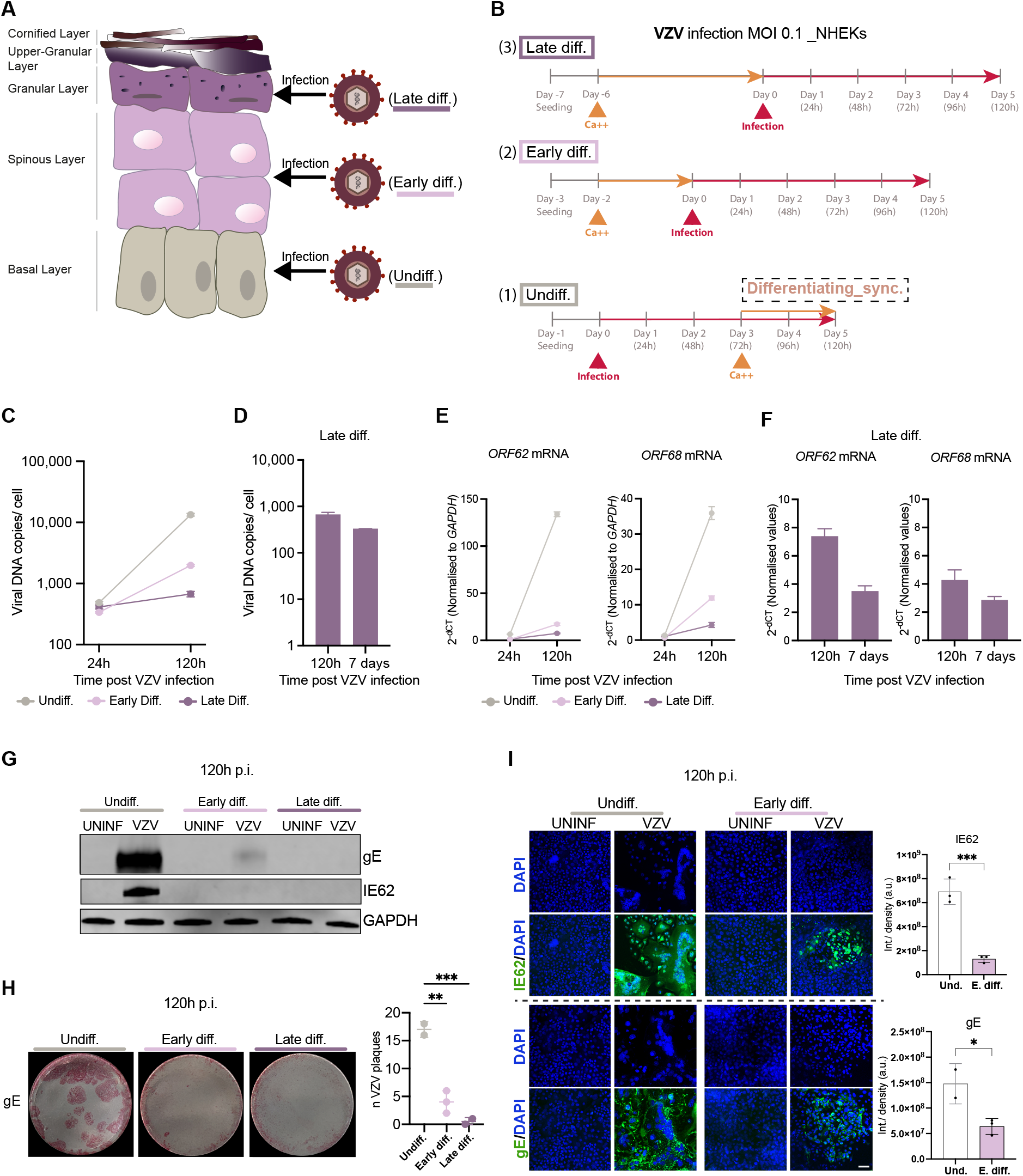
Influence of epidermal terminal differentiation on VZV replication and life cycle. **A**. Schematic of epidermal differentiation and the differentiation stages at which viral infections were implemented; undifferentiated (Undiff.), early differentiated (Early diff.) and late differentiated (Late diff.). **B**. Schematic of plan of infection with VZV at MOI 0.1 of Undiff., Early diff. and Late diff. NHEKs. “Undiff.”, “Early diff.” and “Late diff.” labelling refers to the state of NHEKs differentiation at the time of infection (day 0). The “Undiff.” condition includes the NHEKs undergoing differentiation by calcium switch after infection (Differentiating_sync.) **C**. Analysis by qPCR of cell-associated VZV genome copy number normalised to number of cells, upon infection with VZV of Undiff., Early diff. and Late diff. NHEKs, and evaluated at 24h and 120h p.i. The data are generated from quantification of the VZV *ORF31* gene (gB) normalised to *RPP30* gene (host internal control) and are reported as the mean of three technical replicates ± SD. **D**. Analysis by qPCR of cell-associated VZV genome copy number normalised to number of cells upon infection with VZV of Late diff. NHEKs, and evaluated at 5 days (120h) and 7 days p.i. The data are generated from quantification of the VZV *ORF31* gene (gB) normalised to *RPP30* gene (host internal control) and are reported as the mean of three technical replicates ± SD. **E**. Analysis by qRT-PCR of *ORF62* (encoding the IE62 protein) and *ORF68* (encoding the gE protein) in VZV infection of Undiff., Early diff. and Late diff. NHEKs, and reported at 24h and 120h p.i. The data are reported as mean of 2^-dCT^ of three technical replicates ± SD, where the normalisation was performed against *GAPDH*. **F**. Analysis by qRT-PCR of *ORF62* and *ORF68* in VZV infection of Late diff. NHEKs, and evaluated at 120h and 7 days p.i. The data are reported as mean of 2^-dCT^ of three technical replicates ± SD, where the normalisation was performed against *GAPDH*. **G**. Western Blotting analysis of gE and IE62 expression at 120h p.i. following infection of Undiff., Early diff. and Late diff. NHEKs with VZV. GAPDH was used as loading control. **H**. Fast red staining of gE at 120h p.i. following infection of Undiff., Early diff. and Late diff. NHEKs with VZV. Graph shows the quantification of number of VZV plaques in at least two fields of view per condition and is reported as mean ± SD. **I**. IE62 and gE (green) immunofluorescence staining of Undiff. and Early diff. NHEKs infected with VZV and analysed at 120h p.i. DAPI (blue) was used for staining of nuclei. The immunofluorescence stainings shown are maximum intensity projections. Graphs show quantification of average integrated density of IE62 and gE signals in at least two fields of view per condition ± SD. Data in **C**., **E**. and **G**. are representative of n=3 independent experiments (for the Undiff. and Early Diff. conditions), data in **I**. are representative of n=2 independent experiments. Statistical significance was evaluated in **H**. by one-way ANOVA with Dunnett’s multiple comparisons test, in **I**. by two-tailed t test. Statistical significance is indicated as ^*^P< 0.05, ^**^P< 0.01, ^***^P< 0.001. Scale bar, 50 μm. Undiff., undifferentiated; Early diff., early differentiated; Late diff., late differentiated; p.i., post infection; NHEKs, normal human epidermal keratinocytes.

All three differentiation states were successfully and equally infected by VZV, as indicated by similar VZV DNA copy numbers detected at 24h post infection (**Figure 1C**), a time point at which full VZV replication has not yet occurred (**Figure S2B** and [24]). Replication in undifferentiated keratinocytes resulted in a 28-fold increase in VZV DNA copy number between 24h and 120h (5 days) post infection. In contrast, VZV copy number increased only 6-fold over the same time period when early differentiated (spinous layer) keratinocytes were infected and was negligible when late differentiated (granular layer) cells were infected (**Figure 1C**). To exclude the possibility that VZV replication in early and late differentiated keratinocytes was delayed rather than reduced, the cells were maintained for an additional two days (7 days of infection in total), with no increase in VZV copy number observed (**Figure 1D**).

To better characterise how VZV life cycle is impacted by the differentiation state of the keratinocytes it infects, we evaluated the expression of immediate early (IE62) and late (gE) proteins in these models. We observed a pattern similar to viral copy numbers; the expression of both genes was increased over time in the undifferentiated condition, while there was little or no increase in the early and late differentiated conditions (**Figure 1E**). Specifically, while infection of the undifferentiated cells caused a steep increase in the expression of both *ORF62* (IE62) and *ORF68* (gE) genes (21-fold and 28-fold increase, respectively) between 24h and 120h, this increase was much lower when early and late differentiated keratinocytes were infected (**Figure 1E)**. Furthermore, *ORF62 and ORF68* gene expression did not increase in the late differentiated condition if the infection was maintained for an additional two days, confirming that when infecting late differentiated keratinocytes VZV life cycle is inhibited, not delayed (**Figure 1F)**. At 120h post infection and consistent with gene expression, both IE62 and gE proteins were highly expressed in infected undifferentiated keratinocytes, with negligible or no expression in the infected early and late differentiated (**Figure 1G, H** and **I**). Immunohistochemistry analyses further highlighted that the formation of viral plaques, which in VZV infection present as multinucleated structures (syncytia), is inhibited when the virus infects early or late differentiated keratinocytes (**Figure 1H** and **I**). IE62 is expected to be expressed both in the nucleus and cytoplasm of cells by 120h post infection, when the virus has fully replicated (**Figure S2B**). Instead IE62 displayed this localisation only in the infection of undifferentiated keratinocytes, but remained nuclear in the infection of already differentiated keratinocytes (**Figure 1I)**. Similarly, the gE protein was confined to perinuclear localisation when already differentiated keratinocytes were infected with VZV (**Figure 1I)**.

These data together indicate that while VZV is able to infect suprabasal differentiated keratinocytes, its replication is restricted and this restriction manifests as inhibited viral gene expression, as well as containment of viral proteins at nuclear and perinuclear locations.

### HSV-1 replication and life cycle are restricted when the virus infects already differentiated keratinocytes, although to a lesser extent compared to VZV

We next investigated HSV-1 infection of NHEKs, using the same three keratinocyte models, but infecting at lower MOI (0.005 instead of 0.1) and maintaining the virus in culture for shorter time compared to VZV (72h instead of 120h), reflecting HSV-1’s faster replication cycle (**Figure 2A**). Unlike VZV, addition of calcium to HSV-1 infected cultures appears to inhibit viral replication as evidenced by the reduction in viral copy numbers following calcium treatment of both HSV-1-infected NHEKs (**Figure S3A**) and primary human dermal fibroblasts (NHDFs) (**Figure S3B**). However, the latter do not undergo calcium-mediated differentiation, indicating that the calcium alone may influence HSV-1 replication. Thus, instead of inducing synchronised differentiation with calcium treatment of infected keratinocytes, we allowed HSV-1-infected basal keratinocytes to undergo spontaneous asynchronous differentiation triggered by intracellular calcium signalling induced following cell-cell contact as the cells reach confluence [19–21]. To reduce heterogeneity in the cultures, we seeded NHEKs at slightly higher density than in the VZV infection model, such that although undifferentiated at the time of infection, they reached confluence and initiated contact differentiation by 72h post infection (**Figure 2A** 1.). The similar increase in *KRT10* gene expression over time in both keratinocytes differentiated through calcium-induced differentiation (synchronised differentiation) and spontaneous contact differentiation (asynchronised differentiation) (**Figure S3C**) validated this system.

**Figure 2:**
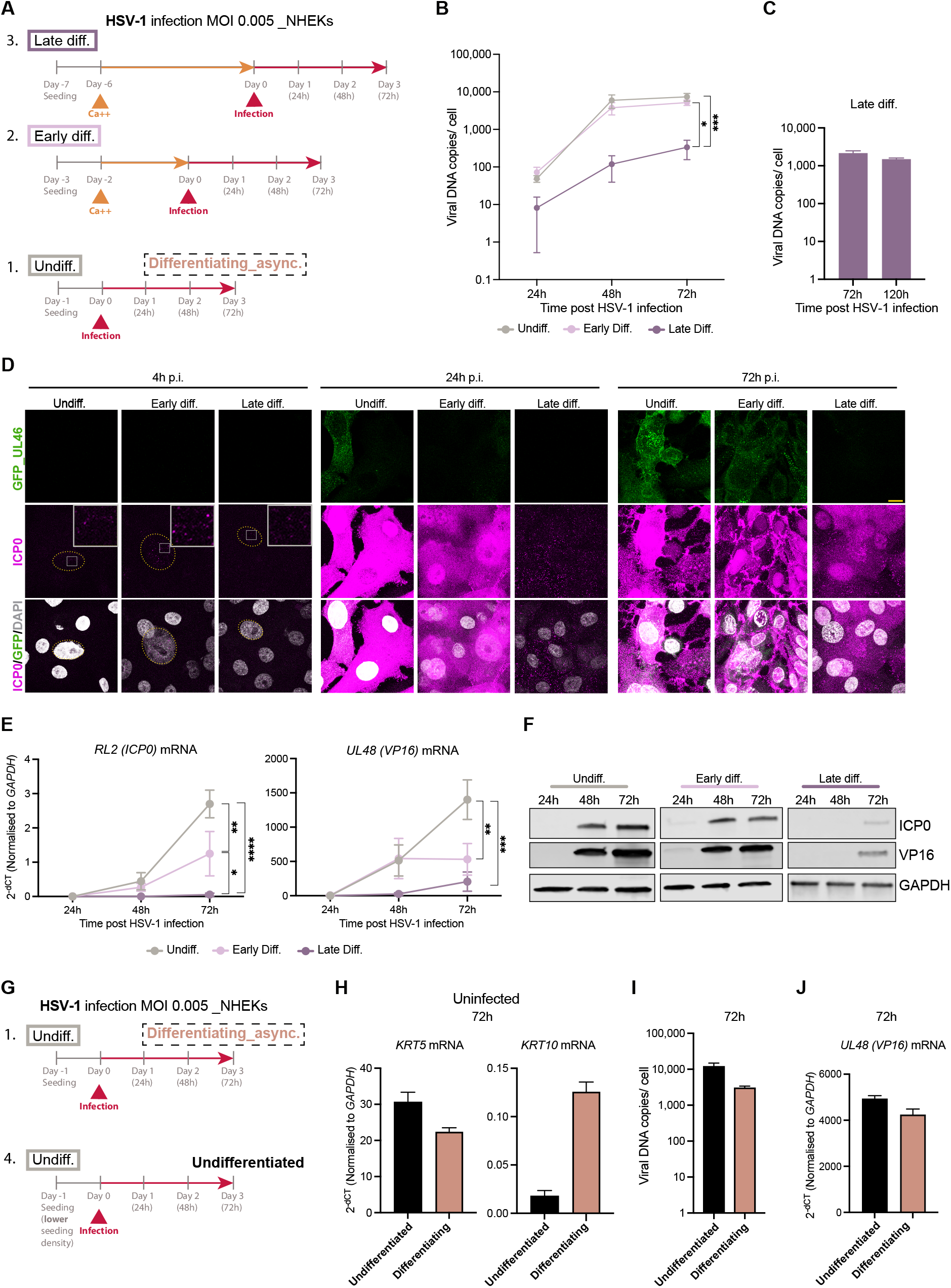
Influence of epidermal terminal differentiation on HSV-1 replication and life cycle. **A**. Schematic of plan of infection with HSV-1 at MOI 0.005 of undifferentiated (Undiff.), early differentiated (Early diff.) and late differentiated (Late diff.) NHEKs. “Undiff.”, “Early diff.” and “Late diff.” labelling refers to the state of NHEKs differentiation at the time of infection (day 0). The “Undiff.” condition includes the NHEKs undergoing spontaneous asynchronous differentiation after infection (Differentiating_async.) **B**. Analysis by qPCR of cell-associated HSV-1 genome copy number normalised to number of cells, upon infection with HSV-1 of Undiff., Early diff. and Late diff. NHEKs, and evaluated at 24h, 48h and 72h p.i. The data are generated from quantification of the HSV-1 *UL27* gene (gB) normalised to *RPP30* gene (host internal control) and are reported as mean of n=3 independent experiments ± SEM. **C**. Analysis by qPCR of cell-associated HSV-1 genome copy number normalised to number of cells, upon infection with HSV-1 of Late diff. NHEKs, and reported at 72h and 120h p.i. The data are generated from quantification of the HSV-1 *UL27* gene (gB) normalised to *RPP30* gene (host internal control) and are reported as the mean of three technical replicates ± SD. **D**. ICP0 (pink) immunofluorescence staining of Undiff., Early diff. and Late diff. NHEKs infected with HSV-1 and analysed at 4h, 24h and 72h p.i. The reported images include GFP (green) signal deriving from the GFP protein tagged to the late HSV-1 tegument protein UL46 and DAPI (grey) staining of nuclei. The immunofluorescence stainings shown are maximum intensity projections. **E**. Analysis by qRT-PCR of *RL2* gene (encoding the ICP0 protein) and *UL48* gene (encoding the VP16 protein) expression in HSV-1 infection of Undiff., Early diff. and Late diff. NHEKs, and reported at 24h, 48h and 72h p.i. The data are reported as mean of 2^-dCT^ of n=3 independent experiments ± SEM, where the normalisation was performed against *GAPDH*. **F**. Western Blotting analysis of ICP0 and VP16 expression at 24h, 48h and 72h p.i. following infection of Undiff., Early diff. and Late diff. NHEKs with HSV-1. GAPDH was used as loading control. Data are representative of n=4 independent experiments (Undiff. condition), n=3 independent experiments (Early diff. condition) and n=2 independent experiments (Late diff. condition). **G**. Schematic of the model of HSV-1 infection implemented in undifferentiated NHEKs, which either remained undifferentiated throughout the infection and until the last time point (72h) (Undifferentiated) or started undergoing spontaneous differentiation during the last time points of infection (Differentiating_async). **H**. Analysis by qRT-PCR of *KRT5* and *KRT10* mRNAs in uninfected NHEKs at 72h conirming the undifferentiated and differentiating status of the keratinocytes. The data are reported as mean of 2^-dCT^ of three technical replicates ± SD, where the normalisation was performed against *GAPDH*. **I**. Keratinocytes were infected when undifferentiated and then either maintained their undifferentiated status throughout the infection (Undifferentiated) or were allowed to spontaneously differentiate by 72h p.i. (Differentiating). Cell-associated HSV-1 genome copy number was analysed at 72h p.i. in both conditions. The data are generated from quantification of the HSV-1 *UL27* gene (gB) normalised to *RPP30* gene (host internal control) and are reported as the mean of three technical replicates ± SD. Similar results were observed in three additional experiments. **J**. Anlysis by qRT-PCR of *UL48* mRNA in HSV-1 infected Undifferentiated and Differentiating NHEKs at 72h p.i. The data are reported as mean of 2^-dCT^ of three technical replicates ± SD, where the normalisation was performed against *GAPDH*. Data are representative of n=2 independent experiments. Statistical significance was evaluated in **B**. and **E**. by 2way ANOVA with Tukey’s multiple comparisons test and indicated as ^*^P< 0.05, ^**^P< 0.01, ^***^P< 0.001, ^****^P< 0.0001. In **D**., dotted lines mark nuclei at 4h p.i. Insets contain zoomed-in areas of the ICP0 signal inside the nuclei at 4h p.i. Scale bar, 10 μm. Undiff., undifferentiated; Early diff., early differentiated; Late diff., late differentiated; p.i., post infection; NHEKs, normal human epidermal keratinocytes.

In contrast to VZV, in which minimal replication was seen, HSV-1 infection of cultures mimicking suprabasal (early differentiated and late differentiated) epithelial layers did result in 72- and 42-fold copy number increases from 24 to 72 hours post infection. Nonetheless, this was reduced compared to the 150-fold increase seen following infection of undifferentiated keratinocytes, suggesting a measure of replication restriction in differentiated cells (**Figure 2B**). At 72h the most significant difference in HSV-1 copy numbers was in the late differentiated condition (**Figure 2B**). To rule out the possibility that the observed phenotype in the late differentiation condition could be caused by delayed HSV-1 replication, we maintained the virus in culture for two additional days (120h-5 days total), with no increase in HSV-1 copy numbers detected (**Figure 2C**).

Because HSV-1 replication cycle is known to be faster than VZV, we investigated whether the difference in the HSV-1 DNA copy numbers 24h post infection (**Figure 2B**) was caused by reduced infectivity of the differentiated conditions, by looking at an earlier time point. At 4h post infection, the punctate nuclear expression of the immediate early protein ICP0 [25,26] and the absence of the late protein UL46 demonstrated that at this time point HSV-1 had only infected the cells and not yet replicated (**Figure 2D**). Conversely, by 24h, replication had occurred as demonstrated by ICP0 widespread expression, including in the keratinocytes’ cytosol, as well as UL46 being expressed (**Figure 2D**). The similar ICP0 protein levels in all three differentiation conditions 4h post infection indicated that HSV-1 was likely to have equally infected keratinocytes at every differentiation state (**Figure 2D**) and that any difference observed at 24h post infection either in expression of viral proteins (**Figure 2D**) or HSV-1 DNA copy numbers (**Figure 2B** and **Figure S4**) was a consequence of different viral replication rates, rather than different infectivity. Furthermore, as with VZV, we observed reduced expression of the viral proteins ICP0 and UL46 in the infected differentiated (early and late differentiated) keratinocytes, with their level of expression being inversely proportionate to the level of differentiation (**Figure 2D**).

To better characterise the HSV-1 life cycle with epidermal differentiation, we investigated the gene and protein expression of both ICP0 and the late protein VP16 between 24h and 72h post infection. Both *RL2* (*ICP0*) and *UL48* (*VP16*) genes were increasingly expressed over time in the infected undifferentiated keratinocytes. In the early differentiated the increase was significantly lower, but higher than in the late differentiated (**Figure 2E**). Similarly, ICP0 and VP16 proteins had the highest expression in the undifferentiated condition, the least in the late differentiated and an intermediate expression in the early differentiated condition (**Figure 2F**).

We previously demonstrated that VZV requires infected keratinocytes to undergo differentiation in order to successfully replicate and mature into infectious virions [13]. To examine whether this is the same in the case of HSV-1 we developed a fourth model of HSV-1 infection where NHEKs were cultured at one third seeding density to guarantee that they remained undifferentiated throughout the 72h course of infection (**Figure 2G** 4.). Over the same time course, while the NHEKs in the first model undergo spontaneous contact differentiation (**Figure 2G** 1._Differentiating_async.), the ones in the fourth model remain undifferentiated (**Figure 2G** 4._Undifferentiated), as validated by high expression of *KRTS* and minimal expression of *KRT10* at the last time point (**Figure 2H**). Cell-associated HSV-1 DNA copy number in the undifferentiated keratinocytes was about 4-fold higher than in the differentiating (**Figure 2I**), although the expression of *UL48* (*VP16*) was not much higher in the undifferentiated compared to the differentiating keratinocytes (**Figure 2J**).

All in all, these data indicate that, although both VZV and HSV-1 are restricted in their replication if infecting suprabasal differentiated keratinocytes, HSV-1 is quite able to productively infect more differentiated cells. However, optimal replication occurs when the virus infects basal undifferentiated keratinocytes.

### VZV and HSV-1 promote a skin blistering phenotype, but use different mechanisms to downregulate the expression of epidermal keratin 10

We previously demonstrated that VZV decreases the expression of keratin 10 (*KRT10*/ K10), a key protein in epidermal terminal differentiation and epidermal barrier function [5,27]. *KRT10* is decreased by VZV at the transcriptional level and the K10 protein is also proteasomally degraded through activation of the kallikrein 6 (KLK6)-MDM2 axis by VZV [13,22]. We also previously reported that this decrease of keratin 10 facilitates VZV replication [22]. To better characterise the dynamics of keratin 10 downregulation in VZV, here we interrogated the expression of keratin 10 early in VZV infection and studied it also in the context of infection of already differentiated keratinocytes in the models described in **Figure 1B**. VZV infection of undifferentiated keratinocytes (which undergo differentiation after infection as per model 1. in **Figure 1B**) resulted in a 92% decrease in *KRT10* gene expression by 24h post infection. By 120h post infection this decrease became 99% (**Figure 3A**) and a similar reduction was observed at the protein level (**Figure 3B** and **C**). Notably, *KRT10* was also decreased at 24 hours when VZV infected early and late differentiated keratinocytes, although with a lower level of reduction (**Figure 3A-C**). No further decrease was observed in *KRT10* when the virus was maintained in culture for two additional days (7 days total of infection) in the late differentiated condition (**Figure 3A**). To examine whether *KRT10* downregulation is related to VZV replication, we used foscarnet treatment, which inhibits viral replication by inhibiting the viral DNA polymerase [28](**Figure S5A**). In infected and differentiating keratinocytes foscarnet treatment had a minimal effect on *KRT10* expression compared to infected cells treated with vehicle alone, indicating that the high *KRT10* gene decrease observed in VZV infection is independent of viral replication (**Figure 3D**).

**Figure 3:**
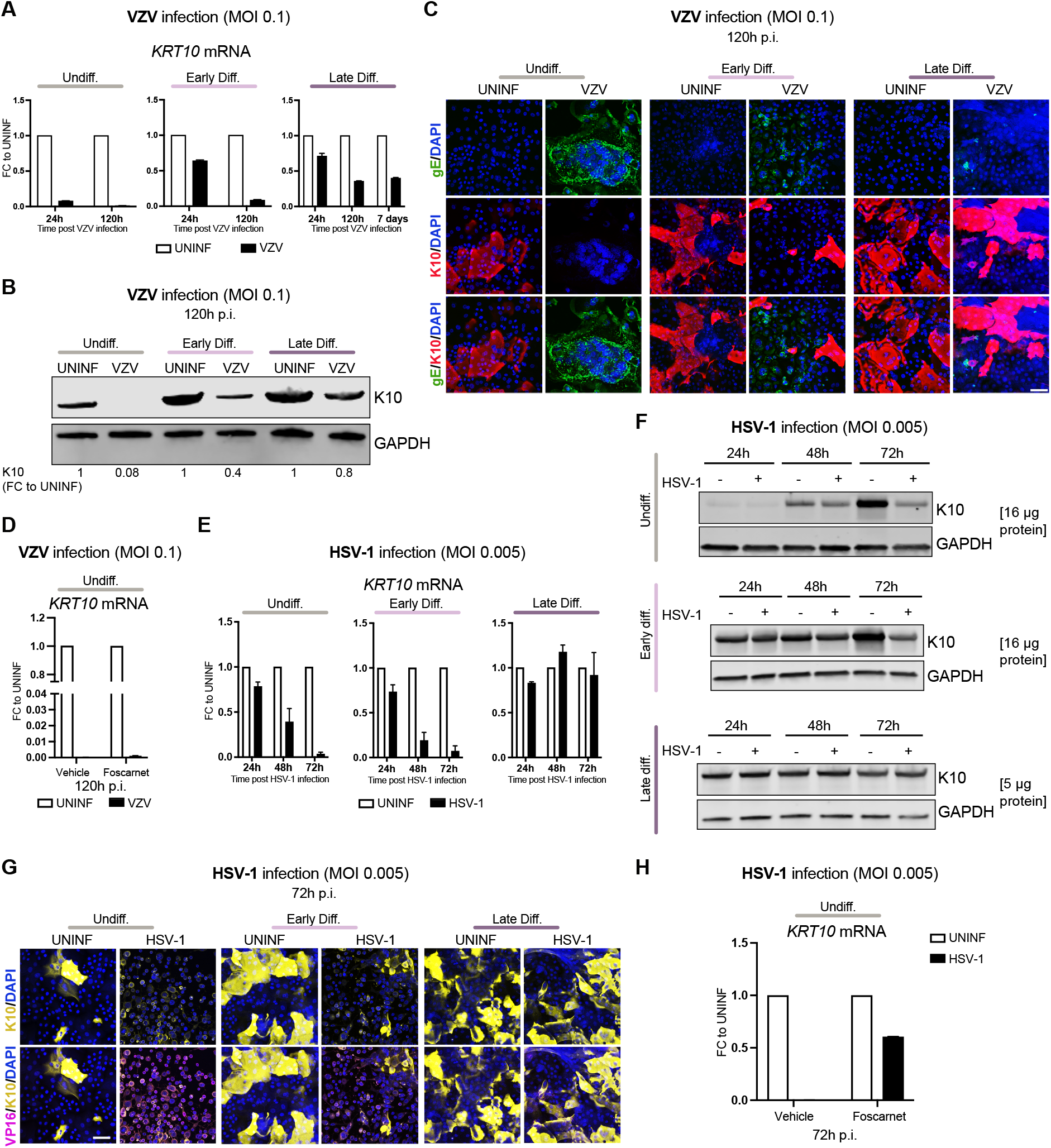
VZV and HSV-1 downregulate keratin 10 expression in infected keratinocytes. **A**. Analysis of qRT-PCR expression of *KRT10* (encoding the K10 protein) in VZV infection (at the MOI of 0.1) of Undiff., Early diff. and Late diff. NHEKs, and reported at 24h, 120h p.i. and at 7 days p.i. (in the Late diff.). The data are reported as fold change (FC) (2^-ddCT^) to each uninfected control from the mean of three technical replicates ± SD, where the normalisation was performed against *GAPDH*. Similar trends were observed in other 2 independent experiments for the Undiff. and Early Diff. conditions and using different MOIs. **B**. Western Blotting analysis of K10 expression at 120h p.i. following infection of Undiff., Early diff. and Late diff. NHEKs with VZV at the MOI of 0.1. GAPDH was used as loading control. Numbers at the bottom of the blots indicate K10 densitometry expressed as fold change to each uninfected control, following normalisation against *GAPDH*. Data are representative of n=3 independent experiments for the Undiff. and Early Diff. conditions. **C**. K10 (red) and gE (green) immunofluorescence staining of Undiff., Early diff. and Late diff. NHEKs infected with VZV at the MOI of 0.1 and analysed at 120h p.i. DAPI (blue) was used for staining of nuclei. Data are representative of n=2 independent experiments for the Undiff. and Early Diff. conditions. **D**. Analysis by qRT-PCR at 120h p.i. of expression of *KRT10* mRNA in VZV infection (at the MOI of 0.1) of Undiff. NHEKs which were treated either with foscarnet or with vehicle alone 48h post infection. The data are reported as fold change (FC) (2^-ddCT^) to each uninfected control, from the mean of three technical replicates ± SD, where the normalisation was performed against *GAPDH*. **E**. Analysis by qRT-PCR of *KRT10* mRNA expression in HSV-1 infection (at the MOI of 0.005) of Undiff., Early diff. and Late diff. NHEKs, and reported at 24h, 48h and 72h p.i. The data are reported as fold change (FC) (2^-ddCT^) to each uninfected control from the mean of n=3 independent experiments ± SEM, where the normalisation was performed against *GAPDH*. **F**. Western Blotting analysis of K10 expression at 24h, 48h and 72h p.i. following infection of Undiff., Early diff. and Late diff. NHEKs with HSV-1 at the MOI of 0.005. GAPDH was used as loading control. Data are representative of n=4 independent experiments for the Undiff. condition, n=3 independent experiments for the Early diff. condition and n=2 independent experiments for the Late diff. condition. In the Undiff. and Early diff. conditions 16μg of proteins were run. In the Late diff. condition 5μg of proteins were run to compensate for the high concentration of K10 in this condition. **G**. K10 (yellow) and VP16 (pink) immunofluorescence staining of Undiff., Early diff. and Late diff. NHEKs infected with HSV-1 at the MOI of 0.005, and analysed at 72h p.i. DAPI (blue) was used for staining of nuclei. **H**. Analysis by qRT-PCR at 72h p.i. of expression of *KRT10* mRNA in HSV-1 infection (at the MOI of 0.005) of Undiff. NHEKs which were treated either with foscarnet or with vehicle alone at the time of infection. The data are reported as fold change (FC) (2^-ddCT^) to each uninfected control, from the mean of three technical replicates ± SD, where the normalisation was performed against *GAPDH*. The data are representative of n=2 independent experiments. Scale bars, 50 μm. Undiff., undifferentiated; Early diff., early differentiated; Late diff., late differentiated; p.i., post infection; UNINF, uninfected.

We next examined *KRT10* gene expression in the three keratinocytes models of HSV-1 infection described in **Figure 2A**. As shown in **Figure 3E**, *KRT10* gene expression decreased progressively over time in HSV-1 infected differentiating keratinocytes (model 1. in **Figure 2A)**, as well as following infection of early differentiated keratinocytes (model 2. in **Figure 2A)**, but not in the late differentiated condition (model 3. in **Figure 2A**), where HSV-1 replication is low (**Figure 2B**), and these findings were confirmed at the protein level (**Figure 3F** and **G**). Contrary to VZV, in HSV-1 infected and differentiating keratinocytes foscarnet-mediated inhibition of viral replication [29] (**Figure SSB**) highly prevented *KRT10* downregulation, with a 40% *KRT10* decrease in keratinocytes treated with foscarnet compared with 99.7% decrease in the infected cells treated with vehicle alone (**Figure 3H**).

Expression of suprabasal desmosome genes, desmocollin-1 (*DSC1*) and desmoglein-1 (*DSG1*), which are downregulated together with *KRT10* by VZV infection [13], were also downregulated by HSV-1 (**Figure S6A** and **B**). Since *DSC1* and *DSG1*, together with *KRT10*, are often downregulated in skin blistering disorders [30,31], the results suggest a similar mechanism for lesion formation in both HSV1 and VZV.

Finally, the downregulation of K10 was confirmed in infections of *ex-vivo* skin tissues with VZV and HSV-1. In VZV, and consistent with what we previously reported [13,22], K10 expression was highly disrupted throughout the tissue and lost in cells positive for the late protein gE (**Figure 4A**). In HSV-1, K10 expression was also disrupted and reduced in the suprabasal cells positive for the late protein UL46 (**Figure 4B**). Notably, while the late VZV protein gE was expressed mainly in the suprabasal layers, the late HSV-1 protein UL46 was mainly expressed in the basal layer and only in some areas suprabasally, confirming that, while expression of late genes and therefore completion of life cycle in VZV is dependent on the infected keratinocytes undergoing differentiation and populating upper epidermal layers [13], HSV-1 can successfully replicate and complete its life cycles in the basal undifferentiated keratinocytes.

**Figure 4:**
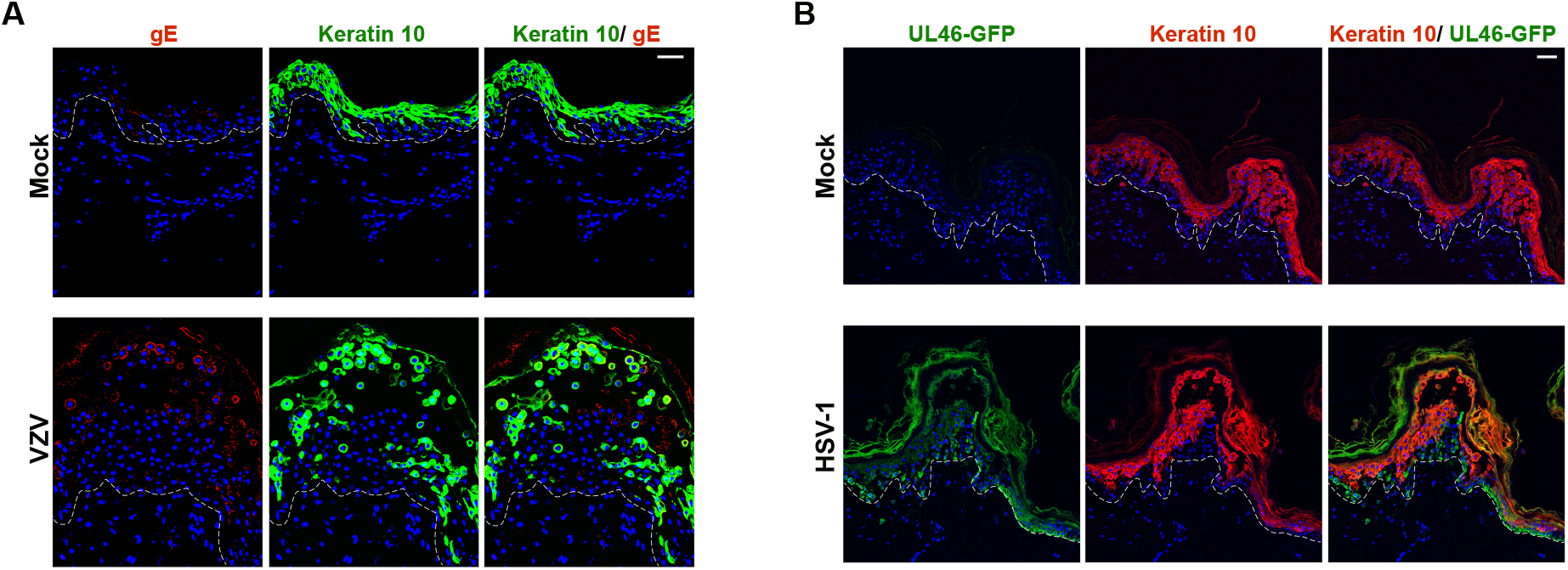
Keratin 10 expression is disrupted in infected cells of *ex vivo* VZV- and HSV-1-infected skin tissues. **A**. Keratin 10 (green) and gE (red) immunofluorescence staining of *ex-vivo* human skin tissues infected with VZV (10^6^ PFU/ml) and analysed 11 days p.i. DAPI (blue) was used for staining of nuclei. **B**. Keratin 10 (red) immunofluorescence staining of *ex-vivo* human skin tissues infected with HSV-1 (10^6^ PFU/ml) and analysed 11 days p.i. GFP signal (green) marks the expression of the GFP-tagged late HSV-1 protein UL46. DAPI (blue) was used for staining of nuclei. The immunofluorescence stainings shown are maximum intensity projections. Dotted lines indicate the epidermal-dermal junctions. Scale bars, 50 μm.

Taken together, these data demonstrate that the downregulation of keratin 10 is a phenotype common to both VZV and HSV-1 infections, however likely achieved through different mechanisms, being this an early event in VZV, while requiring viral replication in HSV-1.

## Discussion

In this study, using the calcium switch model to produce keratinocyte cultures representative of distinct epidermal strata, we show that while VZV replication is severely restricted when suprabasal keratinocytes are infected, HSV-1 is able to replicate following infection of both basal undifferentiated and suprabasal differentiated keratinocytes, although preferentially in the former. These findings provide some measure of clarification for previous observations on HSV-1 replication in skin. Conflicting results have been obtained from infection through microtrauma of skin organotypics or *ex vivo* skin explant models of stratified epithelia, with some authors stating that HSV-1 replication requires infection of basal undifferentiated cells [16–18], while others suggest that infection of the suprabasal spinous layer is optimal [14]. While these 3D models recapitulate epidermal biology, identifying into which of the layers the viral inoculum has been delivered is notoriously difficult. Instead, using host gene expression patterns to define individual epidermal strata, we have been able to confirm the preferential replication of HSV-1 in basal keratinocytes while providing support for its more promiscuous replication in skin compared with VZV.

The calcium switch model also allows us to interrogate more precisely potential host-viral interactions occurring in epidermal infection. While previous observations that VZV late gene expression requires keratinocyte differentiation are confirmed [13], our new data show nuclear retention of viral proteins when suprabasal layers are directly infected (Figure 1I). One explanation could be the loss/ reduction during differentiation of host proteins involved in promoting nuclear export of viral proteins or their cytoplasmic transport. Because epidermal differentiation is a combination of dynamic and spatio-temporal regulated events, several factors could be controlling this. Our data indicate that VZV life cycle, similarly to another family of skin-tropic viruses, the human papillomaviruses [32], follows the progression of epidermal differentiation; VZV initially likely requires some host factors espressed in basal undifferentiated cells to initiate specific phases of its life cycle with subsequent expression of suprabasal proteins to complete it. It is possible that where direct infection of differentiated keratinocytes occurs, some of the host co-factors necessary to VZV replication have been lost thus restricting replicaton. The pattern of nuclear and perinuclear retention of IE62 and gE points to the possible loss in suprabasal layers of host proteins used by VZV in nuclear-cytoplasmic transport. Alternatively, new proteins expressed in the suprabasal cells could restrict virus replication. However, the fact that IE62 and gE expression and protein localisation are not restricted and viral load increases when basal cells infected with VZV undergo differentiation [13], makes this less likely.

In contrast, while HSV-1 replication is most productive in undifferentiated cells, its replication does not appear to be tightly linked to keratinocyte differentiation and indeed replication even diminishes as differentiation occurs (Figure 2I). Unlike VZV, we see no evidence of nuclear retention of immediate early or late proteins when suprabasal layers are directly infected, with evidence of cytoplasmic spread of both even when, overall, their expression is reduced. This suggests differences in how HSV-1 interacts with keratinocyte differentiation compared to VZV. Since we see no change in the localisation of HSV-1 proteins’ staining in cells with differentiation, it is possible that it is reduction in cell-to-cell spread rather than in nuclear-to-cytoplasm spread that restricts HSV-1 suprabasally. One explanation could be a reduced access to HSV-1 cell entry receptors. However, the main HSV-1 keratinocyte receptor, nectin-1 [14,18,31], is expressed in all epidermal layers [18] and in fact its expression appears to increase with differentiation (Tommasi, unpublished data and [33,34]). Thus, if nectin-1 is responsible for this restriction, it may be that the physical access to it is prevented, for example because of a redistribution of the receptor from the cell surface. In contrast, in the case of VZV, two of the three main keratinocytes receptors, the mannose 6 phosphate receptor (M6PR) and the heparan sulphate proteoglycan (HSPG), are expressed mainly in basal undifferentiated keratinocytes ([35,36]), while the IDE (insulin degrading enzyme) receptor is expressed mainly in suprabasal keratinocytes [33,34]. Taken together the results suggest that VZV and HSV-1 use different strategies to replicate in skin and provide pointers as to future experiments that might be used to dissect these out.

Both VZV and HSV-1 are able to downregulate the expression of keratin 10, as well as other suprabasal proteins, resulting in patterns of protein loss analogous to that seen in many genetic and acquired blistering disorders [30]. In VZV infection this occurs almost immediately and does not require viral replication, as indicated by reduced keratin 10 levels at 24 hours before full replication has occurred, in all epidermal strata including in the most differentiated granular layer (where viral replication is minimal), and no change to *KRT10* expression caused by foscarnet-mediated inhibition of replication. This indicates that the downregulation of keratin 10 is likely an early event which might be caused by viral proteins present in the VZV virion. In contrast, in HSV-1, keratin 10 decrease is progressive over time, proportional to the amount of HSV-1 replication, and is reduced when HSV-1 replication is blocked by foscarnet treatment. This suggests a mechanism mediated by proteins that are not present in the virion but rather generated during the course of viral replication. We previously showed that, in addition to transcriptional downregulation of *KRT10*, VZV is also able to promote keratin 10 proteasomal degradation through activation of the KLK6-MDM2 axis, with evidence that this also induces activation of autophagy pathways [22]. Investigations as to whether HSV-1 also employs this second mechanism are underway.

In summary, by employing an approach that enables examination of viral replication at specific epidermal differentiation states, this study identifies similarities and differences in the interplay of VZV/ HSV-1 with epidermal differentiation. Identification of specific epidermal factors responsible for either restriction or facilitation of viral replication in the skin bears potential for novel antiviral strategies, and, by targeting the skin, holds promise for topical applications.

## Materials and methods

### Cell cultures

All cells used in this study were cultured at 37°C in 5% CO2 and regularly tested for mycoplasma contamination using the mycoplasma PCR detection kit (Merck). Primary normal human epidermal keratinocytes (NHEKs) isolated from juvenile foreskin and pooled from different donors were purchased from Promocell or Thermo Fisher Scientific and grown as previously described [22,37]. Briefly, NHEKs were grown on dishes coated with type 1 rat tail collagen (Sigma-Aldrich) and in a 1:1 mixture of keratinocyte SFM medium and medium 154, supplemented with human keratinocyte growth supplement (HKGS) and 1% antibiotics/antimycotics (Thermo Fisher Scientific). In all the experiments the NHEKs used were ≤ 5 passages old.

The three differentiation conditions described in Figure 1A and B (and Figure 2A) were generated with the following set up:

For the Undiff. condition, NHEKs were seeded at a density of 100,000/120,000 (or 150,000 in the case of HSV-1 infection) per well of a 12-well dish.

For the Early diff. condition, the NHEKs were seeded at the density of 250,000 (or 300,000 in the case of HSV-1 infection) per well of a 12-well dish. After one day they were calcium switched for two days with calcium at the concentration of 2.4 mM.

For the Late diff. condition, the NHEKs were seeded at the density of 250,000 (or 300,000 in the case of HSV-1 infection) per well of a 12-well dish. After one day they were calcium switched for six days with calcium at the concentration of 1.2 mM.

Normal human dermal fibroblasts (NHDFs) (Thermo Fisher Scientific) were grown in DMEM medium (Thermo Fisher Scientific) containing 10% FBS and 1% antibiotics/antimycotics (Thermo Fisher Scientific).

### Viruses and viral infections

For infections with VZV, the pOka strain (kind gift from Prof. Paul Kinchington, University of Pittsburgh) was used. The three conditions, “Undiff”, “Early diff.” and “Late diff.” were infected with mitomycin C-treated pOka (day 0) at the MOI of 0.1 and then left in infection for five days. Immediately after infection, cells were maintained at 37°C for 1 hour and then transferred to 34°C. For model 1., the NHEKs were induced to differentiate as previously described in the context of VZV infection, by calcium switch at day 3, with calcium at the concentration of 2.4mM for two days [22,24].

For infections with HSV-1, the SC16 strain and the GFP-tagged SC16 strain (kind gift from Prof. Gillian Elliott, University of Surrey) were used. Some of the experiments of infection of differentiated keratinocytes were repeated with the HSV-1 KOS strain (kind gift from Prof. Matthew Reeves, UCL) and showed equivalent results (data not shown). For infections by HSV-1, NHEKs were infected with HSV-1 at MOI of 0.005, unless otherwise specified in the data, and the same set up as VZV was used, but with different NHEKs starting seeding density (see previous paragraph) and for shorter length of infection, 72h total. In all conditions, one hour after infection the medium containing the virus was replaced with normal medium. For model 1., differentiation was achieved by contact mediated differentiation. Finally, for model 4. described in Figure 2G, NHEKs starting seeding density was 50,000 cells per well of a 12-well dish.

All the viruses’ cultures used in this study were regularly tested for mycoplasma contamination using the mycoplasma PCR detection kit (Merck).

### Skin tissues and viral infections

Human abdominal skin tissues from plastic surgery at Royal Free Hospital were collected with written informed consent under the Research Ethics Committee approval number 24/SW/0096. The tissues were first cut into 0.7×0.7 cm squares and then grown at the air-liquid interface as previously described [22,37] in KGM medium (Lonza) for 12 days. After one day in culture, the tissues were infected by scarification. Briefly, a 27-gauge needle was used to carry out scarification of the epidermis, which was followed by addition of medium containing either 10^6^ PFU/ml of VZV or HSV-1, or medium alone for mock control. After 11 days of infection, the tissues were fixed in formalin, followed by paraffin embedding.

### Quantitative PCR (qPCR)

Genomic DNA was extracted using the AllPrep DNA/RNA Kit (Qiagen) and quantified with Qubit 1x dsDNA Assay Kit (Invitrogen). For each qPCR reaction 5μl of DNA was added to a master mix containing specific primers and probes (at 100 nM concentration) and the QuantiNova Probe PCR Master Mix and QN ROX Reference Dye (QuantiNova Probe PCR Kit; Qiagen). The reactions were subject to 1 cycle of 2 min at 95°C, followed by 45 cycles, each with 5 seconds at 95°C and 10 seconds at 60°C. The sequences of primers and probes used were as reported in literature: for HSV-1 gB (*UL27* gene) [38], for VZV gB (*ORF31* gene) [39], for *RPP30* [40]. DNA copy numbers were extracted from a standard curve based on dilution series of standards whose concentration we previously validated by digital PCR (plasmids containing specific target regions on the viral and human *RPP30* genes were purchased from IDT). *RPP30* was used as internal host control and viral DNA copy numbers were normalised to *RPP30* copy numbers (*RPP30* is present in two copies within human cells) to obtain viral DNA copies per cell.

### Quantitative reverse-transcription PCR (q-RTPCR)

Total RNA was extracted from cells using the RNeasy Plus Mini Kit (Qiagen) or the AllPrep DNA/RNA Kit (Qiagen) and quantified with Qubit RNA XR Assay kit (ThermoFisher Scientific), according to the manufacturer’s instructions. cDNA was synthesised from 1μg of RNA, using the High-Capacity cDNA Reverse Transcription Kit (Thermo Fisher Scientific), according to standard protocols. For qPCR reactions, TaqMan Fast Advanced MasterMix (ThermoFisher Scientific) was used and each reaction was run at 95°C for 20 seconds, followed by 40 cycles at 95°C for 3 seconds and 60°C for 30 seconds. The primers and probes used for host genes were purchased from Thermofisher Scientific, cat n 4448489 (VIC dye): GAPDH (ASSAY ID Hs02786624_g1) and cat n 4331182 (FAM dye): KRT10 (assay ID Hs00166289_m1), KRT1 (assay ID Hs00196158_m1), KRT5 (assay ID Hs00361185_m1), LOR (assay ID Hs01894962_s1), FLG (assay ID Hs00856927_g1), LCE3D (assay ID Hs00754375_s1), DSC1 (assay ID Hs00245189_m1), DSG1 (assay ID Hs00355084_m1). For viral genes, custom primers and probes were purchased from Thermo Fisher Scientific (cat n 4331348, FAM dye) and the sequences are as reported in literature: for VZV *ORF62* and *ORF68* [41], for HSV-1 *RL2* (ICP0) and *UL48* (VP16) [42]. Each qPCR reaction contained one FAM primer and GAPDH-VIC primer for normalisation to *GAPDH*. mRNA levels were analysed as 2^-dCT^ (CT values of the gene of interest were normalised to *GAPDH* CT values) or as 2^-ddCT^, where normalised values were reported as relative values to a condition of interest.

### Western Blotting

Cells were lysed in 5% SDS lysis buffer. Proteins were quantified with Qubit™ Protein BR Assay (Thermo Fisher Scientific) and an amount between 10 and 20μg (unless otherwise specified) of proteins was run in each experiment. The lysates were prepared by mixing with 2x Laemmli Sample Buffer (Bio-Rad) containing 0.2 M DTT, heated for 5 min at 95 °C and run on 4–20% Tris-Glycine gel (Bio-Rad). The transfer of proteins on PVDF membrane (Bio-Rad) was performed by using Bio-Rad Trans-Blot Turbo. The PVDF membranes were then blocked in Odyssey® Blocking Buffer (LICORbio) for 30 minutes at room temperature, then incubated overnight at 4°C with primary antibody solution in Odyssey® Blocking Buffer with 0.1% Tween-20 and containing primary antibodies against the target protein and GAPDH as the loading control. After washes, the membranes were then incubated for 1 hour with secondary antibodies at the concentration of 1:15000 in Odyssey® Blocking Buffer containing 0.1% Tween-20 and 0.01% SDS. The membranes were finally imaged on a LI-COR Odyssey CLx and the fluorescence was analysed by Image Studio™ Software (LICORbio). Primary antibodies: K10 (1: 1000; Biolegend, 905404), GAPDH (1: 1000; MILLIPORE, MAB374), HSV-1 ICP0 (1: 500; Santa-Cruz, sc-53070), HSV-1 VP16 (1: 1000; abcam, ab110226), VZV IE62 (1:1000; Abcam, ab212015), VZV gE (1:500; Santa Cruz, sc-56995). Secondary antibodies: goat anti-rabbit IRDye® 680RD (LICOR, 926-68071), goat anti-mouse IRDye® 800CW (LICOR, 926-32210), goat anti-rabbit IRDye® 800CW (LICOR, 926-32211), goat anti-mouse IRDye® 680RD (LICOR, 926-68070).

### Immunohistochemistry for plaques assay (Fast red staining)

Fast Red staining of VZV plaques was performed as previously described in [13] and using ready-to-use Fast Red TR/Naphthol AS-MX tablets. For staining of VZV plaques the gE antibody (Santa Cruz, sc-56995) at the concentration of 1:50 was used.

### Immunofluorescence

NHEKs were grown on collagen-coated coverslips and infected as described in previous paragraphs. At the end of the experiment, cells were fixed and permeabilised with a solution of 4% PFA and 0.2 % Triton-X100. Blocking was performed for 30 min with a solution containing 0.4% fish skin gelatin (Merck) and 0.2% Triton X-100, followed by 1 hour incubation with primary antibodies and finally 1 hour incubation with secondary antibodies at the concentration of 1:500. For stainings of skin tissues, the blocking step was preceded by deparaffinisation in xylene and antigen retrieval in 0.01 M Na Citrate (pH 6).

Both cells and tissues were mounted in Prolong Gold Anti-fade Reagent with DAPI (Life Technologies) and imaged on a Zeiss LSM710 microscope. Imaging analyses were performed in Fiji-ImageJ [43].

#### Primary antibodies

VZV IE62 (1:200; abcam, ab212015), HSV-1 ICP0 (1:100; Santa Cruz, sc-53070), HSV-1 VP16 (1:100, abcam, ab110226), K10 (1: 500; Invitrogen, MA106319) or (1:500; Biolegend, 905404), VZV gE (1:50; Santa Cruz, sc-56995), GFP (1:500, antibodies, A290). Secondary antibodies: goat anti-mouse Alexa Fluor 488 (Life Technologies, A11029), goat anti-mouse Alexa Fluor 555 (Life Technologies, A21424), goat anti-mouse Alexa Fluor 647 (Life Technologies, A21235), goat anti-rabbit Alexa Fluor 555 (Life Technologies, A32732), goat anti-rabbit Alexa Fluor 594 (Life Technologies, A11012), goat anti-rabbit Alexa Fluor 488 (Life Technologies, A11008).

### Drug treatment

Foscarnet (or vehicle alone) was added at the concentration of 50 ug/ml to the NHEKs’ medium at the time of HSV1 infection and 48h post VZV infection and in both cases maintained in the medium until the end of the experiment.

## Abbreviations

VZV: Varicella-zoster virus
HSV-1: herpes simplex virus 1
NHEKs: normal human epidermal keratinocytes
NHDFs: normal human dermal fibroblasts
*KRT10*: keratin 10 mRNA
K10: keratin 10 protein
*KRT1*: keratin 1 mRNA
*KRTS*: keratin 5 mRNA
*KRT14*: keratin 14 mRNA
*IVL*: involucrin mRNA
*LOR*: loricrin mRNA
*FLG*: filaggrin mRNA
*LCE3D*: late cornified envelope 3D mRNA
*DSC1*: desmocollin-1 mRNA
*DSG1*: desmoglein-1 mRNA
Undif: undifferentiated
Early diff.: early differentiated
Late diff.: late differentiated
sync.: synchronised
async.: synchronised.

## Declarations

### Ethics approval

Human skin tissue was collected with written informed consent under the Research Ethics Committee approval number 24/SW/0096.

### Conflict of interests statement

The authors state no conflict of interest.

### Funding

This work was funded by the John Black Charitable Foundation and the Rosetrees Trust. CT was supported by the John Black Charitable Foundation with initial funding from National Institutes of Health Grant R01AI158510. AD was supported by the John Black Charitable Foundation. GK was supported by the Rosetrees Trust. JB receives funding from the NIHR UCLH/UCL BRC.

### Authors’ contribution

**CT**: Conceptualization, data curation, formal analysis, investigation, validation, visualization, methodology, funding acquisition, supervision, project administration, writing-original draft, writing-review and editing. **GK**: Data curation, formal analysis, investigation, validation, visualization, methodology. **AL**: Investigation, validation, visualization, methodology. **AD**: Investigation, methodology. **OO**: Investigation **OETM**: Formal analysis, validation, methodology. **AM**: Investigation **JB**: Conceptualization, data curation, methodology, funding acquisition, supervision, project administration, writing-original draft, writing-review and editing.

## Acknowledgements

Microscopy was performed at the Light Microscopy Core Facility, UCL GOS Institute of Child Health supported by the NIHR GOSH BRC award.

The histopathology work was supported by the NIHR GOSH BRC. The views expressed are those of the authors and not necessarily those of the NHS, the NIHR or the Department of Health.

## Notes

### Competing Interest Statement

The authors have declared no competing interest.

### Summary of Updates

Some additional experimental data has been included in Figure 3 and Figure S5 and S6.

