## Supplementary material for "Comparative analysis of varicella-zoster virus and herpes simplex virus 1 interaction with epidermal terminal differentiation in primary human keratinocytes models of differentiation"

### Supplementary Figures:

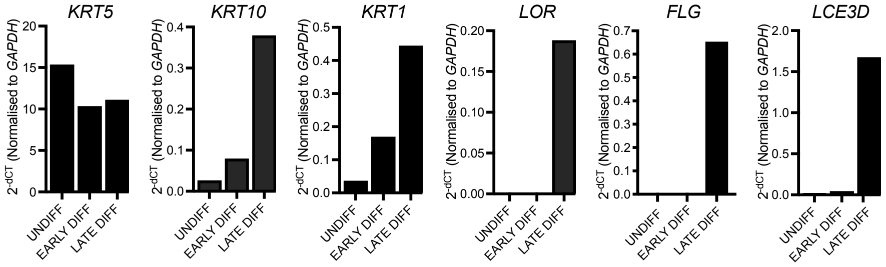


**Figure S1 : Expression of epidermal differentiation markers in the three differentiation states evaluated in the study.**

Analysis by qRT-PCR of expression of epidermal differentiation markers *KRT5* (encoding the keratin 5 protein), *KRT10* (encoding the keratin 10 protein), *KRT1* (encoding the keratin 1 protein), *LOR* (encoding the loricrin protein), *FLG* (encoding the filaggrin protein), and *LCE3D* (encoding the late cornified envelope 3D protein) in Undiff., Early diff. and Late diff. NHEKs. The data are reported as mean of $2^{-dCT}$ of three technical replicates, where the normalisation was performed against *GAPDH*. Undiff., undifferentiated; Early diff., early differentiated; Late diff., late differentiated.

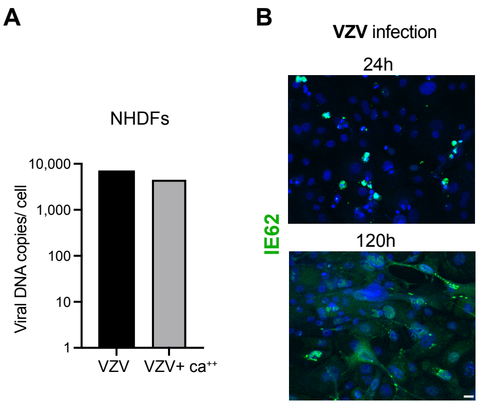


**Figure S2 : Effect of calcium on VZV replication. Evaluation of VZV IE62 expression over time of infection.**

**A.** Analysis by qPCR of VZV genome copy number normalised to number of cells, upon infection with VZV at the MOI of 0.1 of normal human dermal fibroblasts (NHDFs) with or without addition of calcium (at concentration of 2.4mM) to the cells' media. **B.** IE62 (green) immunofluorescence staining of NHEKs infected with VZV at the MOI of 0.1 and analysed at 24h and 120h p.i. DAPI (blue) was used for staining of nuclei. The nuclear expression of IE62 at 24h post infection indicates that the virus has not yet replicated at this time point, whereas widespread expression of IE62, including in cells' cytoplasm, at 120h post infection indicates full replication.

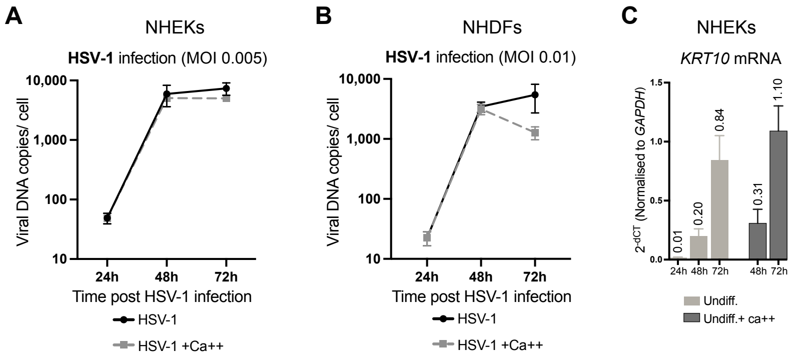


**Figure S3 : Effect of calcium on HSV-1 replication and keratinocytes differentiation.**

**A.** Analysis by qPCR of HSV-1 genome copy number normalised to number of cells, upon infection with HSV-1 at the MOI of 0.005 of NHEKs with or without addition of calcium (at concentration of 2.4mM) to the cells' media, and evaluated 24h, 48h, 72h p.i. The data are reported as mean± SEM of n=3 independent experiments. **B.** Analysis by qPCR of HSV-1 genome copy number normalised to number of cells, upon infection with HSV-1 at the MOI of 0.01 of NHDFs with or without addition of calcium (at concentration of 2.4mM) to the cells' media, and evaluated 24h, 48h, 72h p.i. The data are reported as mean± SEM of n=2 independent experiments. **C.** Analysis by qRT-PCR of *KRT10* (encoding the keratin 10 protein) expression over time (24h, 48h, 72h) in undifferentiated NHEKs with (Undiff. +ca++) or without (Undiff.) addition of calcium (at concentration of 2.4mM) 24h post seeding. The data are reported as mean± SEM of $2^{-dCT}$ of n=3 independent experiments, where the normalisation was performed against *GAPDH*.

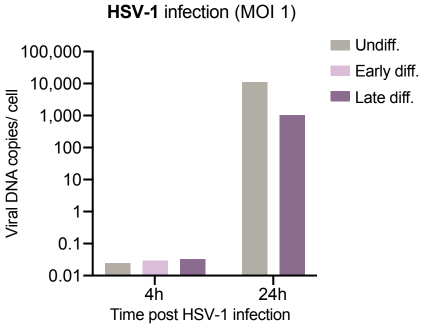


**Figure S4 : HSV-1 DNA copy numbers at 4h and 24h p.i. in infection of the three differentiation conditions.**

Analysis by qPCR of cell-associated HSV-1 genome copy number normalised to number of cells, upon infection with HSV-1 at the MOI of 1 of Undiff., Early diff. and Late diff. NHEKs and analysed at 4h and 24h p.i. Data are the mean of three technical replicates. Undiff., undifferentiated; Early diff., early differentiated; Late diff., late differentiated.

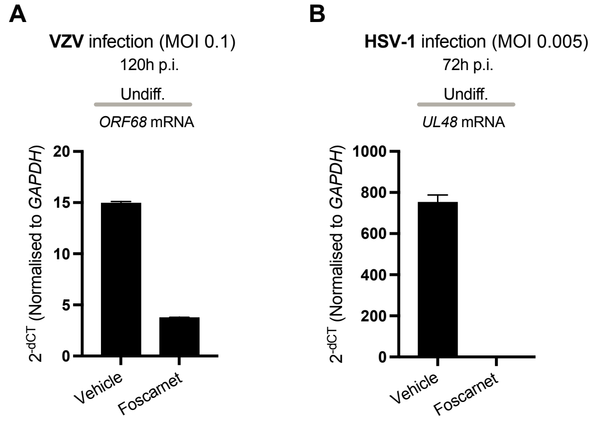


**Figure S5 : Foscarnet inhibits the expression of the late VZV gene *ORF68* and the late HSV-1 gene *UL48* in infection of keratinocytes.**

**A.** Analysis by qRT-PCR of expression of *ORF68* gene at 120h p.i. in VZV infection at MOI of 0.1 of Undiff. NHEKs (as per model in **Figure 1 B.** 1.) with or without addition of foscarnet 48h post infection. The data are reported as mean of $2^{-dCT}$ of three technical replicates ± SD, where the normalisation was performed against *GAPDH*. **B.** Analysis by qRT-PCR of expression of *UL48* gene at 72h p.i. in HSV-1 infection at MOI of 0.005 of Undiff. NHEKs (as per model in **Figure 2 A.** 1.) with or without addition of foscarnet at the time of infection. The data are reported as mean of $2^{-dCT}$ of three technical replicates ± SD, where the normalisation was performed against *GAPDH*. Data are representative of n=2 independent experiments.

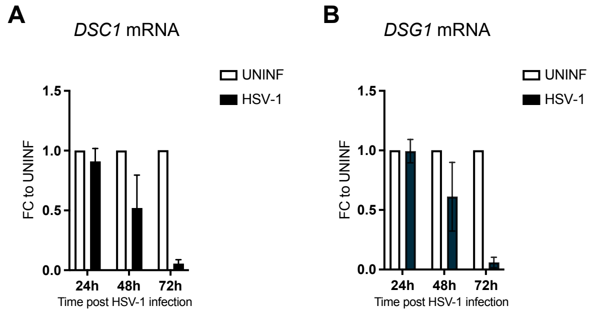


**Figure S6: HSV-1 downregulates *DSC1* and *DSG1*.**

**A.** Analysis by qRT-PCR of *DSC1* (encoding the desmocollin-1 protein) expression in HSV-1 infection (at the MOI of 0.005) of Undiff. NHEKs (as per model in **Figure 2 A.** 1.), and reported at 24h, 48h and 72h p.i. The data are reported as fold change (FC) ($2^{-ddCT}$) to each uninfected control from the mean of n=3 independent experiments ± SEM, where the normalisation was performed against *GAPDH*. **B.** Analysis by qRT-PCR of *DSG1* (encoding the desmoglein-1 protein) expression in HSV-1 infection (at the MOI of 0.005) of Undiff. NHEKs (as per model in **Figure 2 A. 1.**), and reported at 24h, 48h and 72h p.i. The data are reported as fold change (FC) ($2^{-ddCT}$) to each uninfected control from the mean of n=3 independent experiments ± SEM, where the normalisation was performed against *GAPDH*.
